# Myostatin gene deletion alters gut microbiota stimulating fast-twitch glycolytic muscle growth

**DOI:** 10.1101/2022.07.24.501334

**Authors:** Zhao-Bo Luo, Shengzhong Han, Xi-Jun Yin, Hongye Liu, Junxia Wang, Meifu Xuan, Chunyun Hao, Danqi Wang, Yize Liu, Shuangyan Chang, Dongxu Li, Kai Gao, Huiling Li, Biaohu Quan, Lin-Hu Quan, Jin-Dan Kang

**Affiliations:** Key Laboratory of Natural Medicines of the Changbai Mountain, Ministry of Education, College of Pharmacy, Yanbian University; Yanji, 133002, China; Department of Animal Science, College of Agricultural, Yanbian University; Yanji, 133002, China; Jilin Provincial Key Laboratory of Transgenic Animal and Embryo Engineering, Yanbian University; Yanji, 133002, China; College of Integration Science, Yanbian University; Yanji, 133002, China

**Author notes:** Correspondence: Lin-Hu Quan, Key Laboratory of Natural Medicines of the Changbai Mountain, Ministry of Education, College of Pharmacy, Yanbian University, Yanji, 133002, China, Email address., Jin-Dan Kang, Department of Animal Science, College of Agricultural, Yanbian University; Yanji, 133002, China. These authors contributed equally (Zhao-Bo Luo, Shengzhong Han, Xi-Jun Yin).

## Abstract

The host genome may influence the composition of the intestinal microbiota, and intestinal microbiota performs an important role in muscle growth and development. Here, we showed that Myostatin (MSTN), a key factor for muscle growth, deletion alters muscularis, plica, and intestinal barrier in pigs. Mice transplanted with *MSTN^−/−^* pig intestinal flora showed increase in the cross-sectional area of myofibers and fast-twitch glycolytic muscle mass. The microbes responsible for the production of short chain fatty acids (SCFAs) were enriched in both *MSTN^−/−^* pigs and recipient mice, and SCFAs levels were elevated in the colon contents. We demonstrated that valeric acid can stimulate type IIb myofiber growth by activation of the Akt/mTOR pathway via GPR43 and improve muscle atrophy induced by dexamethasone. This is the first study to identify the *MSTN* gene-gut microbiota-SCFA axis and its regulatory role in fast-twitch glycolytic muscle growth.

## Introduction

The decline in muscle mass is a considerable health problem that deteriorates the quality of life and increases disease occurrence and mortality (Newman et al., 2006; Srikanthan et al., 2014). For instance, a decrease in muscle mass contributes to the onset of various diseases, such as sarcopenia, obesity, diabetes, and cancer. Myostatin (MSTN), a transforming growth factor β family member, is among the major regulators of skeletal muscle growth and development (Chen et al., 2021). Substantial muscle hypertrophy was observed in *MSTN* mutant animals and humans (McPherron et al., 1997; Ceccobelli et al., 2022; McPherron and Lee, 1997; Kambadur et al., 1997; Mosher et al., 2007; Kang et al., 2017; Schuelke et al., 2004). Recently various MSTN inhibitors, including monoclonal antibodies, have been tested in clinical trials to treat muscle disorders, such as sarcopenia and cancer-associated cachexia (Kim et al., 2021; Cho et al., 2022). Notably, *MSTN* is not only expressed in skeletal muscles, but also in smooth muscles including the intestine, to participate in various metabolic processes (Sundaresan et al., 2008; Verzola et al., 2017; Esposito et al., 2020; Kovanecz et al., 2017). Previous studies have shown that *MSTN* mutation can alter the composition of intestinal flora in pigs (Pei et al., 2021). However, the interaction between the gut microbiota reshaped by *MSTN* deletion and the host is unclear.

Genetic variation can reshape the structure of the gut microbiota. Mutation in human *SLC30A2* leads to reduced intestinal zinc transport and increased *Clostridiales* and *Bacteroidales* abundance, causing mucosal inflammation and intestinal dysfunction (Kelleher et al., 2022). Moreover, the gut *GLUT1* gene deletion altered the abundances of *Barnesiella intestinis* and *Faecalibaculum rodentium*, promoted fat accumulation, and impaired sugar tolerance (He et al., 2022). These results suggest that host genes can influence the gut microbiota, thereby regulating physiological processes. Intestinal structural changes, such as in intestinal length, epithelial thickness, and surface area by surgery, could affect intestinal function and microbial composition (Seganfredo et al., 2017; Nicoletti et al., 2017; Agus et al., 2018). Barrier defects were accompanied by major changes in the fecal microbiota and a significantly decreased abundance of *Akkermansia muciniphila*, increaseing the vulnerability to gastrointestinal disorders (Sovran et al., 2019).

The intestinal microbiota plays a crucial role in muscle growth and development. For example, urease gene-rich microbes, *Alistipes* and *Veillonella* respectively maintain muscle mass in hibernating animals by promoting urea nitrogen salvage (Regan et al., 2022) and metabolize lactic acid to provide energy for skeletal muscles for long periods of exercise and increase endurance in runners (Scheiman et al., 2019). Short chain fatty acids (SCFAs) are gut microbiota-derived metabolites that are involved in maintaining the integrity of the intestinal mucosa, improving glucose and lipid metabolism, controlling energy expenditure, and regulating the immune system and inflammatory responses (Agus et al., 2021; Besten et al., 2013). SCFAs are absorbed in gut lumen and mediate host metabolic responses in various organs, including skeletal muscle (Frampton et al., 2020). SCFAs play a vital role in skeletal muscle mass maintenance (Lv et al., 2021; Chen et al., 2022), and are involved in the regulation of lipid and glucose metabolism primarily through G protein-coupled receptors (GPRs), such as GPR41, GPR43, and GPR109 (Stoddart et al., 2008; Hul et al., 2019).

Skeletal myofibers exhibit remarkable diversity and plasticity in energy metabolism and contractile functions. Slow-twitch muscles are rich in mitochondria and have high oxidative capacity, whereas fast-twitch muscles generate ATP primarily through glycolysis (Schiaffino et al., 2011; Bassel-Duby et al., 2006). Aging and muscle atrophy result in a gradual decline in muscle mass and strength accompanied by a higher proportion of type I myofibers, leading to muscle weakness due to the preferential loss and atrophy of fast-twitch glycolytic type IIb myofibers (Akasaki et al., 2014; Haber et al., 1992; Faulkner et al., 2007; Kirkendall et al., 1998). Type IIb myofibers are larger in size and more glycolytic and generate high contractile force, but have poorer resistance to fatigue than type I myofibers (Schiaffino et al., 2011). The activation of Akt/mTOR was confirmed to promote the transition from oxidized to glycolytic myofiber types by elevating the levels of glycolytic proteins HK2, PFK1, and PKM2 (Meng et al., 2013; Izumiya et al., 2008; Verbrugge et al., 2020).

MSTN can affect the growth and function of skeletal muscles. This study aimed to investigate whether the intestinal flora remodeled by *MSTN* deletion is involved in the regulation of skeletal muscle growth. Because pigs are highly similar to humans in many aspects such as physiology, disease progression and organ structure (Swindle et al., 2012), we used *MSTN*^−/−^ pigs to investigate the effects of *MSTN* deletion on intestinal structure and the relationship between intestinal microbiota and skeletal muscle growth and function and to explore the underlying mechanisms involved in the regulation of muscle growth by the *MSTN* gene–gut microbiota–skeletal muscle axis.

## Results

### MSTN deletion stimulates muscle hypertrophy and alters intestinal structure and composition of gut microbiota in pigs

We used *MSTN^−/−^* pigs with 2 and 4 bp deletions in the two alleles of the *MSTN* gene (Figure 1-figure supplement 1A). They were generated using the TALEN genome editing technique (Kang et al., 2017). We found that those pigs had higher skeletal muscle mass and myofiber CSA but lost MSTN expression and reduced phosphorylation of smad2/3 in skeletal muscles (Figure 1A-C). The protein expression of myosin heavy chain (MyHC) type IIb, MyoD and glycolytic enzymes HK2, PFK1 and PKM2 were significantly increased in skeletal muscle (Figure 1C, D). The expression of MSTN was not detected in intestine, whereas that of smooth muscle proteins α-SMA and calponin-1 was increased (Figure 1E). We also observed an increase in muscularis thickness and plica length of intestinal and upregulated expression of tight junction-related genes *ZO-1* and *Occludin* (Figure 1F, G). These findings indicate that *MSTN* knockout leads to changes in intestinal structure.

**Figure 1.**
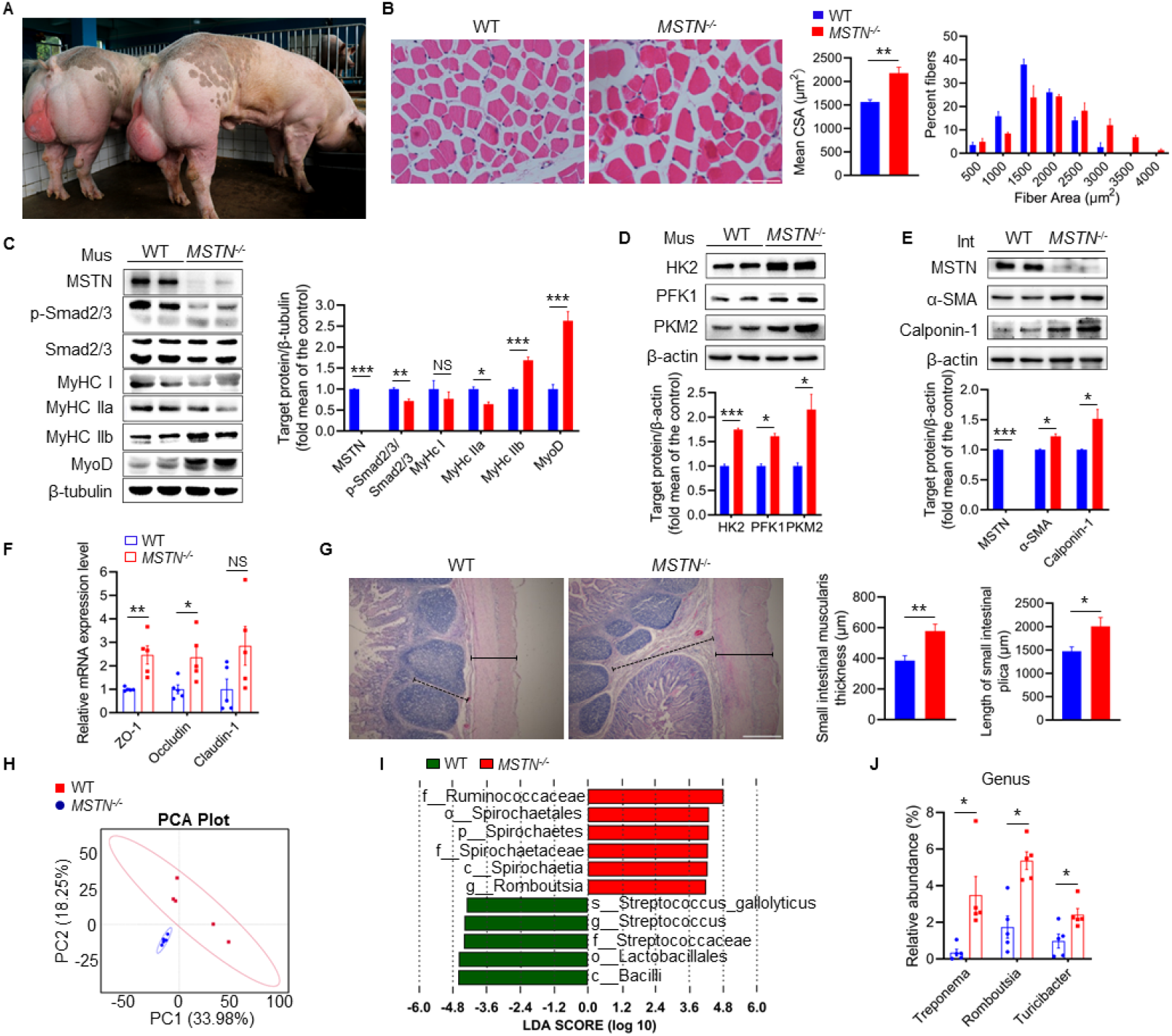
*MSTN* deletion stimulates muscle hypertrophy and alters intestinal structure and composition of gut microbiota in pigs (n=5). (**A** and **B**) Representative images of *MSTN^−/−^* pigs and hematoxylin eosin staining longissimus dorsi. Magnification is 200×. Scale bar, 100 μm. *MSTN^−/−^* pigs showed skeletal muscle hypertrophy and significantly increased the muscle fiber area. (**C**) Relative to WT pigs, *MSTN^−/−^* pigs showed no expression of MSTN, downregulate phosphorylation of Smad2/3 and MyHC IIa, and upregulate MyHC IIb and MyoD in longissimus dorsi (Mus). (**D**) *MSTN^−/−^* pigs showed increased glycolysis enzymes HK2, PFK1 and PKM2 in longissimus dorsi (Mus). (**E**) The protein expression of MSTN was not detected in intestine (Int) while the α-SMA and Calponin-1 were increased in *MSTN^−/−^* pigs compared with the WT pigs. (**F**) Relative expression of tight junction genes *ZO-1* and *Occludin* were enhanced in small intestine of *MSTN^−/−^* pigs. (**G**) Hematoxylin eosin staining of intestinal morphology. The dotted line indicates the length of the plica and the solid line indicates the thickness of muscularis. Magnification is 40×. Scale bar, 500 μm. *MSTN^−/−^* pigs showed an increase of muscularis thickness and plica length in small intestine. (**H**) Plots shown were generated using the weighted version of the Unifrac-based PCA. (**I**) Discriminative taxa determined by LEfSe between two groups (log10 LDA>4.8). (**J**) Comparison proportion of genus levels in feces detected by pyrosequencing analysis showed *Treponema*, *Romboutsia*, and *Turicibacter* were increased in *MSTN^−/−^* pigs. Statistical analysis is performed using Student’s *t-test* between WT and *MSTN^−/−^* pigs. Data are means ± SEM. \**p* < 0.05; \*\**p* < 0.01; \*\*\**p* < 0.001; NS, not statistically significant.

Because host genotypes and phenotypes in various mammals interact with the gut microbiota (Kreznar et al., 2017), we speculated that *MSTN* deletion could affect the composition of the gut microbiota by altering intestinal structure. Thus, fecal samples from *MSTN^−/−^* and wild-type (WT) pigs were examined to determine the diversity and abundance of gut microbiota using 16s rRNA-based microbiota analysis. The alpha-diversity values showed that the ACE in *MSTN^−/−^* pigs are significantly lower than that in WT pigs; however, Chao 1, Shannon, and Simpson indexes were no significant difference (Figure 1-figure supplement 1B-E). These results suggest that *MSTN* deficiency can lead to a decrease in the abundance of intestinal flora. The composition structure of the gut microbiota, as analyzed by PCA, showed that the two groups can be clearly differentiated (Figure 1H). LEfSe analysis confirmed a significant difference at the genus level in *Romboutsia* (Figure 1I). In addition, *Treponema*, *Romboutsia*, and *Turicibacter* were significantly increased at the genus level (Figure 1J). Notably, these altered genera are involved in SCFAs production (Kreznar et al., 2017; Li et al., 2019b; Li et al., 2021; Li et al., 2019c; Bian et al., 2020). These results verify that *MSTN* deficiency can alter the intestinal structure while promoting the growth of microbes related to SCFAs production.

### Gut microbiota reshaped by *MSTN* gene deletion promotes fast-twitch glycolytic muscle growth

To determine the effect of the *MSTN*-deleted altered intestinal flora on skeletal muscle, we transplanted fecal microbes from *MSTN^−/−^* pigs and WT pigs into mice. Mice translated with WT pig feces were named WT-M, and those with *MSTN^−/−^* pig feces were named KO-M. After eight weeks of normal chow feeding, KO-M had a higher muscle mass than to WT-M, especially an enlarged gastrocnemius (GA) muscle (Figure 2A). The GA mass, but not that of the soleus (SOL) or extensor digitorum longus (EDL), was significantly enhanced in KO-M than in WT-M (Figure 2B). However, there was no significant difference in food intake, physical activity, energy intake, or absorbed energy between the two mice groups (Figure 2-figure supplement 2A-E).

**Figure 2.**
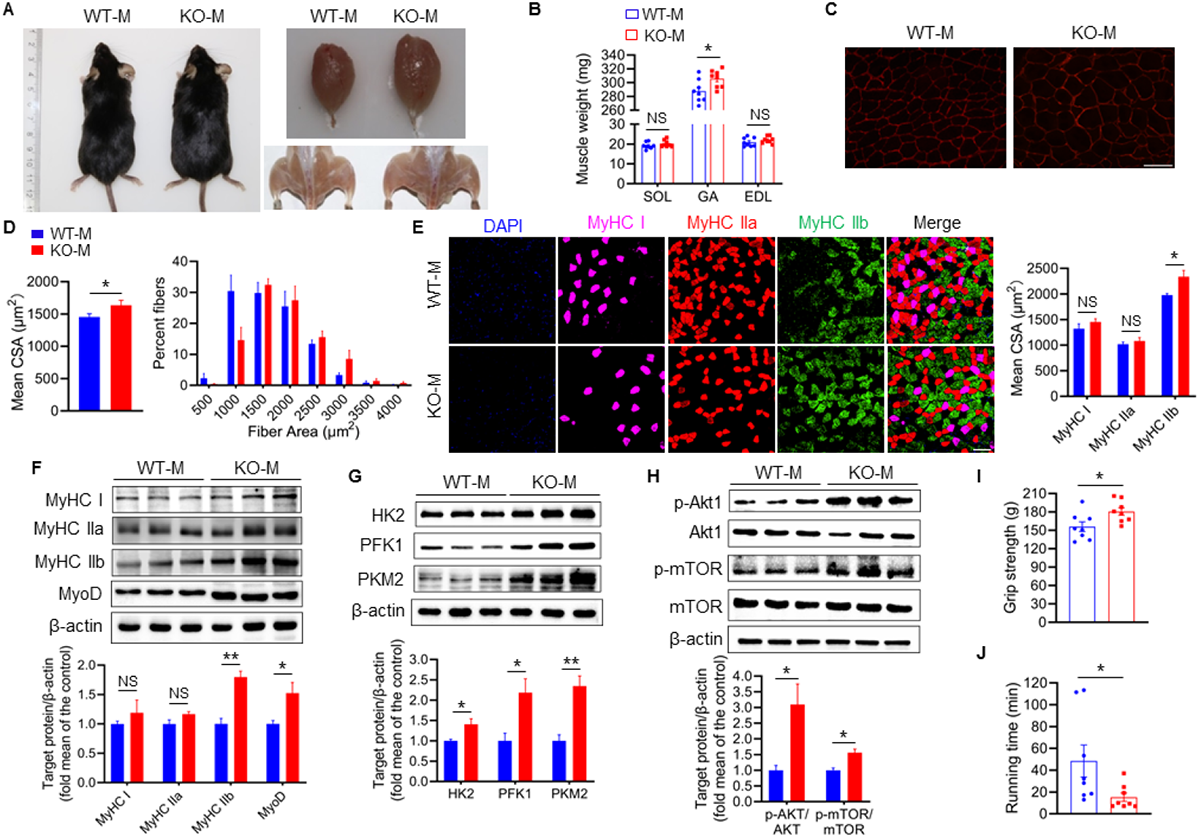
Mice fecal microbiota transplantation from *MSTN* deletion pigs induces type IIb myofiber growth. Mice were treated with porcine fecal microbiota for eight weeks by daily oral gavage after combined antibiotics treatment for a week. WT-M, WT pigs fecal microbiota-received mice (n=8); KO-M, *MSTN^−/−^* pigs fecal microbiota-received mice (n=8). (**A**) Representative images of gross appearance and GA of WT-M and KO-M. (**B**) GA mass was increased in KO-M while SOL and EDL were not different between WT-M and KO-M. (**C**) Representative images of GA sections stained with laminin. Magnification is 400×. Scale bar, 50 μm. (**D**) Quantification analysis of myofiber CSA showed that KO-M was larger than WT-M. (**E**) Representative images of GA sections stained with MyHC I (pink), IIa (red), IIb (green) antibodies and nucleuses were stained with DAPI (blue). Magnification is 200×. Scale bar, 100 μm. Quantification of myofiber displayed MyHC IIb CSA were in GA. (**G**) The expression of glycolysis enzymes HK2, PFK1 and PKM2 were increased in KO-M GA. (**H**) The Akt/mTOR pathway was activated in KO-M GA. (**I** and **J**) Grip strength was enhanced while running time was reduced in KO-M compared with WT-M. Statistical analysis is performed using Student’s *t-test* between WT-M and KO-M groups. Data are means ± SEM. \**p* < 0.05; \*\**p* < 0.01; NS, not statistically significant.

Quantitative analysis of fiber size of GA muscle revealed that the CSA of fiber is significantly hypertrophic in KO-M, and that the distribution of fiber sizes in KO-M clearly shifted toward larger fibers (Figure 2C, D). As shown in Figure 2E, the CSA of type IIb myofibers in KO-M was markedly higher than that in WT-M. Correspondingly, the levels of proteins MyHC IIb and MyoD and those of glycolytic enzymes HK2, PFK1 and PKM2 were significantly increased in the GA muscle of KO-M, whereas the levels of MyHC I and IIa were not significantly different (Figure 2F, G). Interestingly, we observed an increase in Akt and mTOR phosphorylation in the skeletal muscle of KO-M (Figure 2H). The Akt/mTOR signaling pathway affects type IIb myofiber hypertrophy (Izumiya et al., 2008; Dutchak et al., 2018), suggesting an explanation for the increased GA mass in KO-M.

We also performed a series of physiological experiments to evaluate the strength and running performance of fecal microbiota transplantation (FMT) mice. Similar to the expression profile of type IIb myofibers, the grip force of KO-M increased compared with that of WT-M (Figure 2I). However, KO-M had a reduced capacity for running (Figure 2J). Owing to an enlargement in type IIb myofibers, a type of fast-twitch glycolytic muscle, which resulted in a higher explosive force and a lower endurance, KO-M had higher grip strength but shorter running time. Collectively, these observations strongly indicate that KO-M have increased CSA of type IIb myofibers and significantly enhanced fast-twitch glycolytic skeletal muscle mass.

### *MSTN^−/−^* pigs FMT alter gut microbiota composition in mice

To investigate the correlation between myofiber hypertrophy and intestinal microbiota in mice, we analyzed their intestinal microorganisms. There were no significant differences in the ACE, Chao 1, Shannon, and Simpson indexes for alpha-diversity (Figure 3-figure supplement 3A-D). Principal coordinates analysis (PCoA) showed that the microbiota composition structure of the two groups is clearly differentiated (Figure 3A). In addition, *Romboutsia* was significantly enriched at the order-, family-, genus levels in KO-M intestinal flora (Figure 3B). The heat map showed that abundance of 22 of the 35 increased genera and 13 decreased genera. *Romboutsia*, which was upregulated in *MSTN^−/−^*pigs, was also upregulated in KO-M (Figure 3C). KO-M was similar to *MSTN^−/−^* pigs, LEfSe analysis showed that *Romboutsia* abundance increased at the genus level (Figure 3D). Functional prediction analysis showed that intestinal microbial functions are concentrated in pathways related to metabolite synthesis (including K05349 and K01952) in KO-M (Figure 3E). These results showed that the mice translated with *MSTN^−/−^* pig feces had increased *Romboutsia* abundance in the intestine.

**Figure 3.**
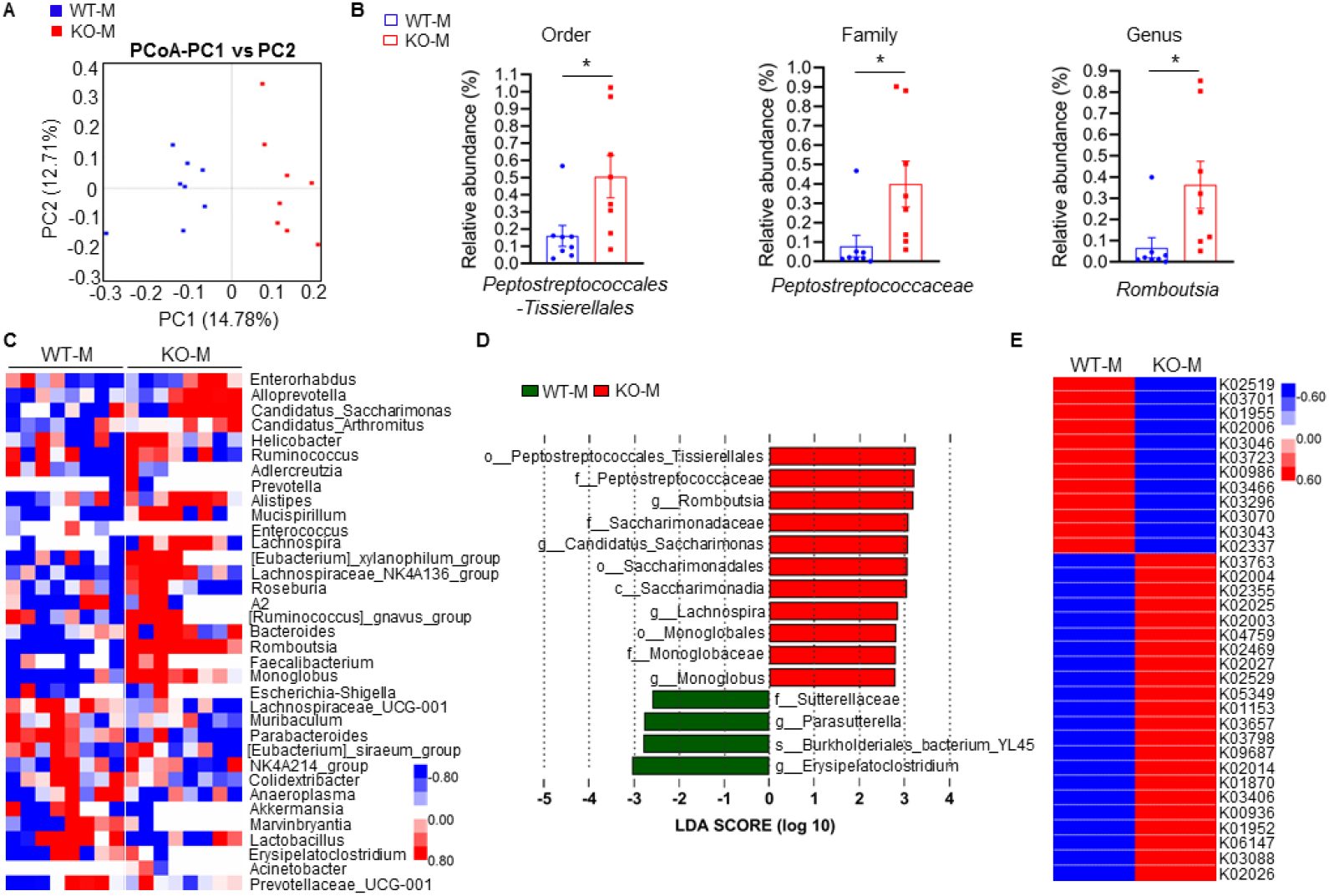
*MSTN*^−/−^ pigs fecal microbiota transplantation alters microbiota composition in mice. Transplanting fecal microbiota of *MSTN^−/−^* pigs and WT pigs separately to mice (n=8). (**A**) Plots shown were generated using the weighted version of the Unifrac-based PCoA. (**B**) Comparison proportion of order, family and genus levels of *Romboutsia* in feces detected by pyrosequencing analysis. (**C**) Heatmap shows the abundance of top 35 microbial genuses levels was significantly altered by WT and *MSTN^−/−^* donor pigs between WT-M and KO-M groups. (**D**) Discriminative taxa determined by LEfSe between two groups (log10 LDA>3.5). (**E**) Functional prediction shows that intestinal microbial functions are concentrated in functional pathways related to metabolite synthesis after fecal microbiota transplantation. Statistical analysis is performed using Student’s *t-test* between WT-M and KO-M groups. Data are means ± SEM. \**p* < 0.05.

### Gut microbes derivative-valeric acid promote myogenic differentiation of myoblasts

As metabolites of the intestinal flora, SCFAs, can affect the growth and function of skeletal muscle (Frampton et al., 2020). The results of our FMT experiments showed that *MSTN* deletion-mediated intestinal microbiota significantly increases skeletal muscle mass and simultaneous enrich *Romboutsia* which can produce SCFAs. Further analysis of fatty acids in the colon contents of mice showed that SCFAs are enriched in KO-M than WT-M; particularly, valeric acid and isobutyric acid were significantly enhanced in KO-M (Figure 4A). However, medium-chain fatty acids (MCFAs) showed no significant differences between the two groups (Figure 4B). Long-chain fatty acids (LCFAs) also showed no difference in WT-M and KO-M overall, although FFA18:2 and FFA16:0 were significantly decreased in KO-M (Figure 4C). The heatmap also confirmed that the differences in SCFAs between WT-M and KO-M (Figure 4D).

**Figure 4.**
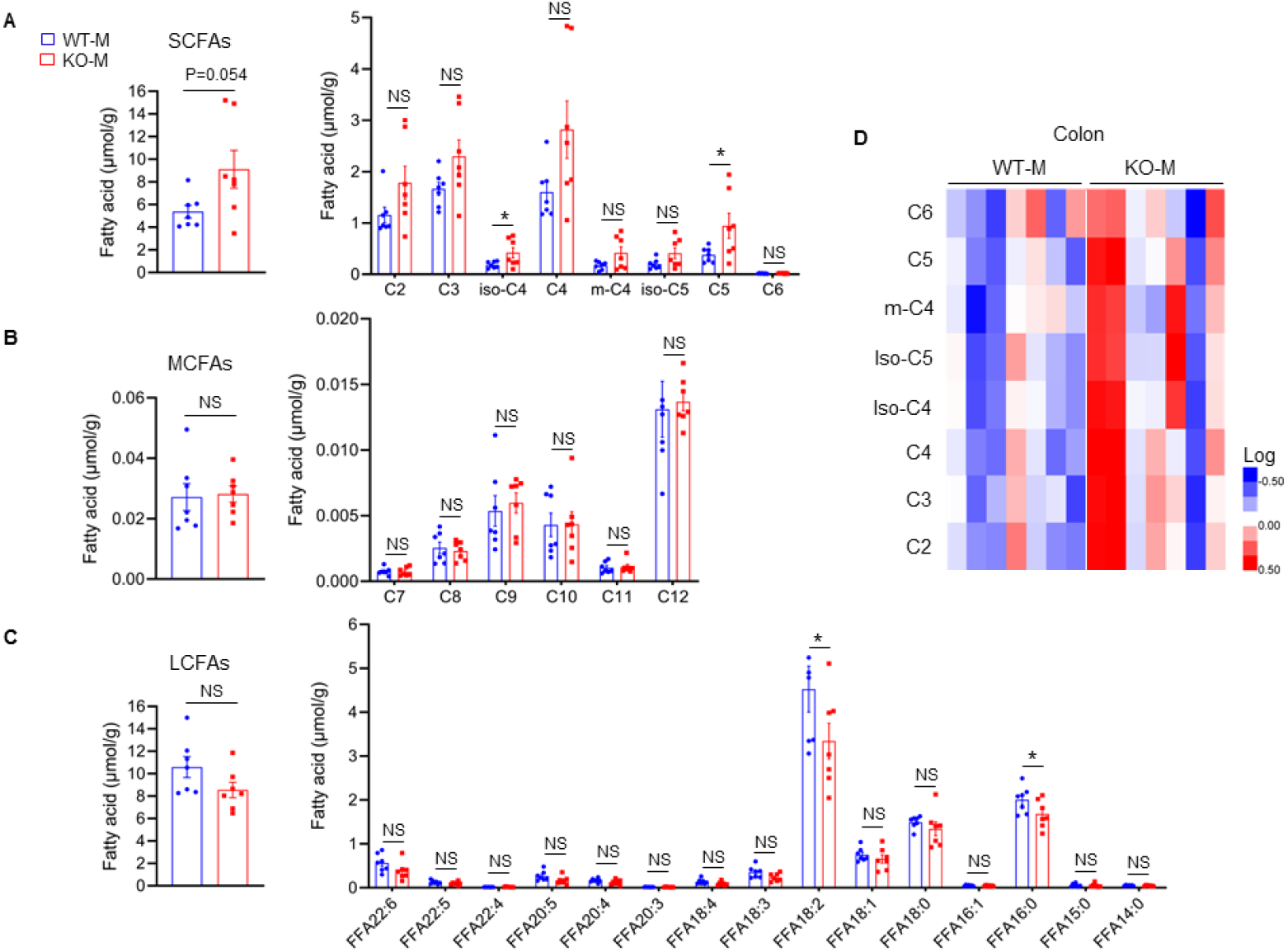
*MSTN^−/−^* pigs fecal microbiota transplantation alters the level of fatty acids in mice (n=7). (**A**) Fecal microbiota transplantation increased colon total SCFAs (particularly valeric acid and isobutyric acid) in KO-M. (**B**) Fecal microbiota transplantation has no effect on MCFAs between WT-M and KO-M. (**C**) Fecal microbiota transplantation decreased colon FFA18:2 and FFA16:0 of LCFAs in KO-M. (**D**) Heatmap showed the difference of SCFAs between WT-M and KO-M. Statistical analysis is performed using Student’s *t-test*. Data are means ± SEM. \**p* < 0.05; NS, not statistically significant.

To assess the influence of upregulated SCFAs on myoblast differentiation, the C2C12 myoblast cell line was treated for 24 h with 5 mM each of valeric acid and isobutyric acid during differentiation. Immunofluorescence staining of MyHC showed that after supplementation of valeric acid, C2C12 myoblasts produced thicker myotubes and notably higher fusion index than the control cells, implying that valeric acid promotes myotube formation (Figure 5A). Valeric acid treatment also improved the expression of MyoD and MyoG and promoted the differentiation of C2C12 myoblasts (Figure 5B). Notably, the phosphorylation levels of Akt and mTOR significantly increased after valeric acid treatment (Figure 5C). However, isobutyric acid treatment did not show such effects and only increased the myotube fusion index. Taken together, these results strongly demonstrate that valeric acid can promote myogenic differentiation of myoblasts.

**Figure 5.**
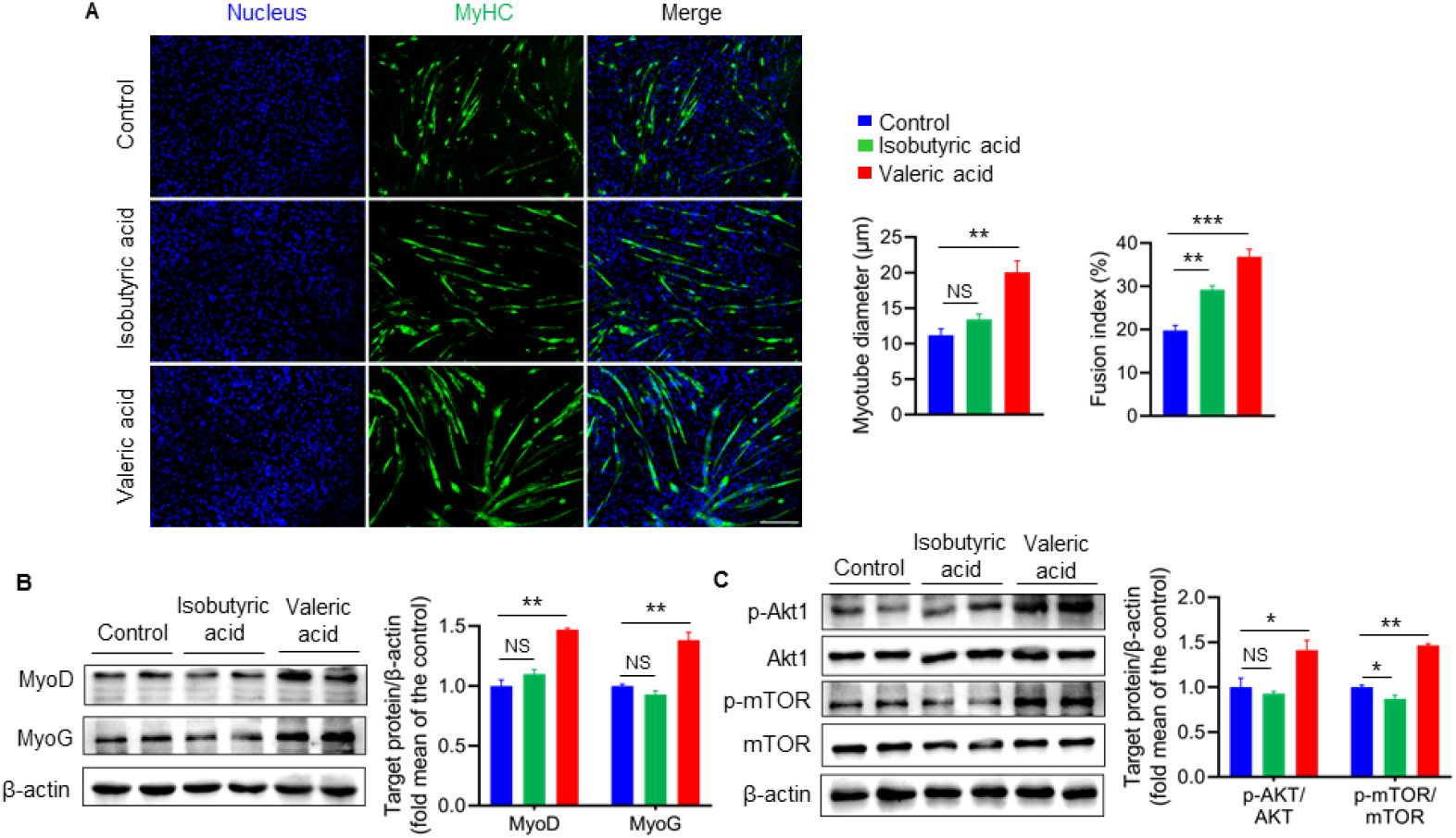
Valeric acid treatment promotes myogenic differentiation of myoblast (n=6). (**A**) Representative images of immunofluorescence stained with a specific antibody to identify MyHC (green) of myotubes and the nucleuses were stained with DAPI (blue). Magnification is 100×. Scale bar, 200 μm. Quantification analysis displayed valeric acid treatment increased the diameter and fusion index of myotube, while isobutyric acid only increased the myotube fusion index. (**B**) Valeric acid treatment increased the expression of MyoD and MyoG in C2C12 myoblasts. (**C**) Valeric acid treatment activated the Akt/mTOR pathway. Statistical analysis is performed using one-way ANOVA. Data are means ± SEM. \**p* < 0.05; \*\**p* < 0.01; \*\*\**p* < 0.001; NS, not statistically significant.

### Valeric acid stimulates type IIb myofibers growth

We further elucidated the effect of valeric acid treatment on the phenotype of skeletal muscles *in vivo*. Mice were administered with valeric acid (100 mg/kg) by daily oral gavage. Valeric acid treatment significantly increased the mass of GA muscle, a fast-twitch glycolytic skeletal muscle, compared with the control (Figure 6A, B). Consistently, following valeric acid treatment, the CSA of the GA muscle was significantly larger, and there was a higher proportion of large myofibers compared with the control (Figure 6C). In valeric acid-treated mice, the protein expression of the MyHC IIb was significantly enhanced, that of MyHC I was decreased, and that MyHC IIa showed no change (Figure 6D). In addition, valeric acid treatment significantly upregulated the levels of glycolysis enzymes of HK2, PFK1, and PKM2 (Figure 6E) and the phosphorylation of Akt and mTOR in the GA muscle (Figure 6F).

**Figure 6.**
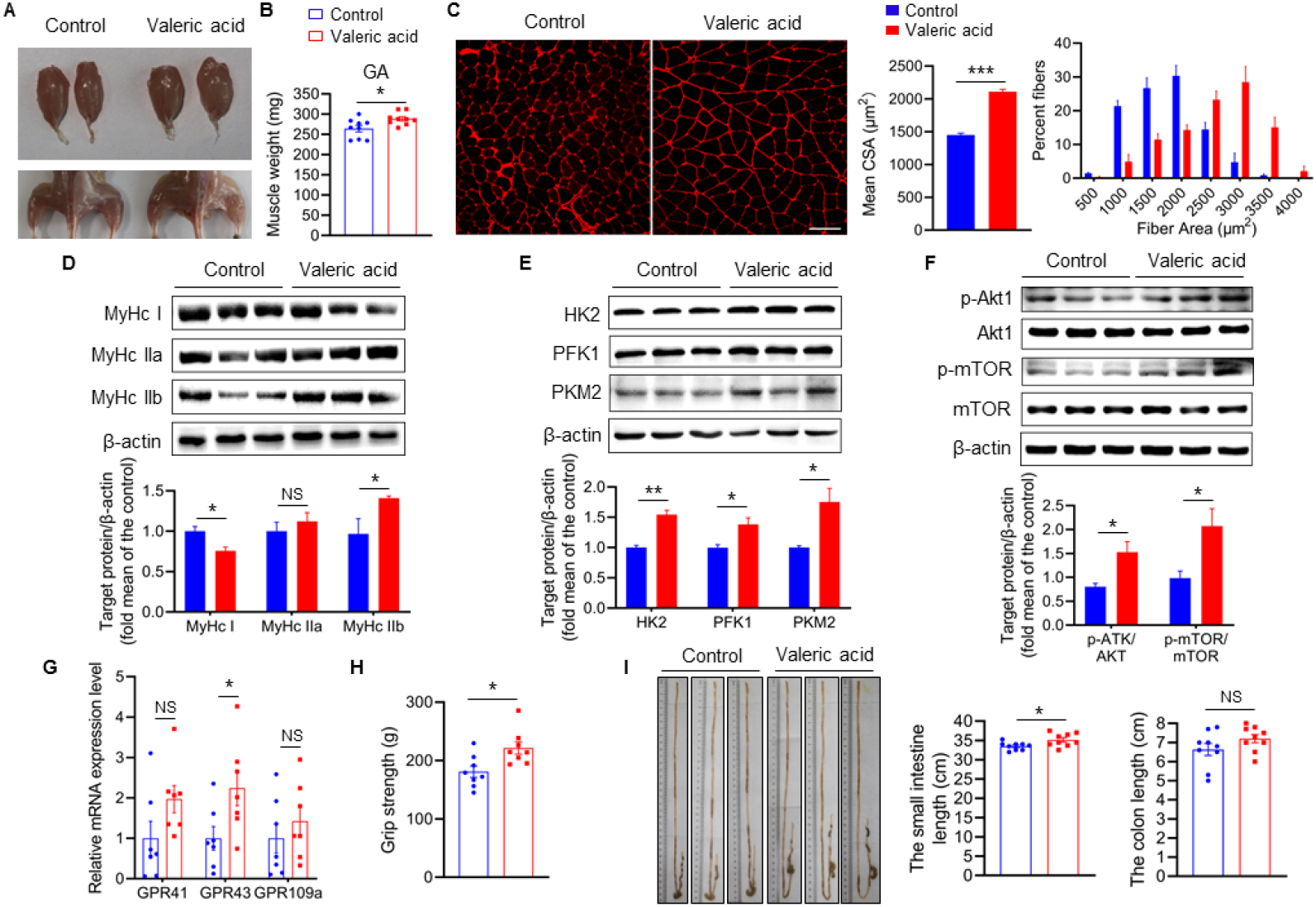
Valeric acid induced type IIb myofiber growth and increased GA mass in mice. Mice were treated with valeric acid (100 mg/kg) for five weeks by daily oral gavage (n=8-9). (**A**) Representative images of gross appearance and GA of control and valeric acid treated mice. (**B**) Valeric acid treatment increased GA mass. (**C**) Representative images of GA sections stained with laminin, showed valeric acid treatment increased CSA of myofiber. Magnification is 200×. Scale bar, 100 μm. Western blot analysis showed that valeric acid treatment increased the levels of (**D**) MyHC IIb, (**E**) glycolysis enzymes HK2, PFK1 and PKM2, and (**F**) activated the Akt/mTOR pathway in GA compared with control mice. (**G**) Real-time PCR analysis indicated that valeric acid treatment enhanced relative mRNA expression of SCFAs receptor *GPR43* in GA. (**H**) Valeric acid treatment improved grip strength. (**I**) Representative images of cecum, small intestine, and colon of mice, showed valeric acid treatment inceresed small intestine length. Statistical analysis is performed using Student’s *t-test*. Data are means ± SEM. \**p* < 0.05; \*\**p* < 0.01; \*\*\**p* < 0.001; NS, not statistically significant.

To explore whether the regulatory pathway, mediated by valeric acid on muscle mass growth would be dependent on fatty acid receptors, we examined the expression of SCFAs receptors in skeletal muscle. Valeric acid increased the mRNA level of *GPR43* in GA muscle, whereas those of *GPR41* and *GPR109a* did not significantly change (Figure 6G). SCFAs could activate the Akt/mTOR pathway through GPR43 (Tang et al., 2022; Brown et al., 2003). These findings suggest that valeric acid, a metabolite of gut microbes, can induce type IIb/glycolytic myofiber growth and enhance GA mass by activating Akt/mTOR signalling through GPR43. Furthermore, valeric acid-treated mice had a greater grip force (Figure 6H). Interestingly, valeric acid treatment increased the length of the small intestine (Figure 6I), but had no effect on food intake, physical activity, energy intake, or absorbed energy in mice (Figure 6-figure supplement 4A-E).

### Valeric acid ameliorates dexamethasone (Dex)-induced skeletal muscle atrophy

Glucocorticoids, such as dexamethasone, are often used to induce muscle atrophy models, and are implicated in protein metabolism in skeletal muscle and are considered as a risk factor for the development of muscle atrophy (Hong et al., 2019; Li et al., 2017). To further explore the role of valeric acid in skeletal muscles, we constructed Dex-induced *in vivo* and *in vitro* muscular atrophy models. Valeric acid administration partially ameliorated skeletal muscle atrophy induced by Dex in mice and reduced the dissolution area with a clear morphology of muscle fiber (Figure 7A). Meanwhile, valeric acid treatment significantly decreased the mRNA and protein levels of muscular dystrophy factors Atrogin-1 and MuRF-1, which were induced by Dex (Figure 7B, C). In C2C12 myoblasts, valeric acid treatment significantly increased myotube diameter and fusion index, and inhibited the expression of atrophy factors, which can improve Dex-induced myotube atrophy (Figure 7D, E). Overall, these finding indicate that valeric acid has a positive effect on Dex-induced muscle atrophy.

**Figure 7.**
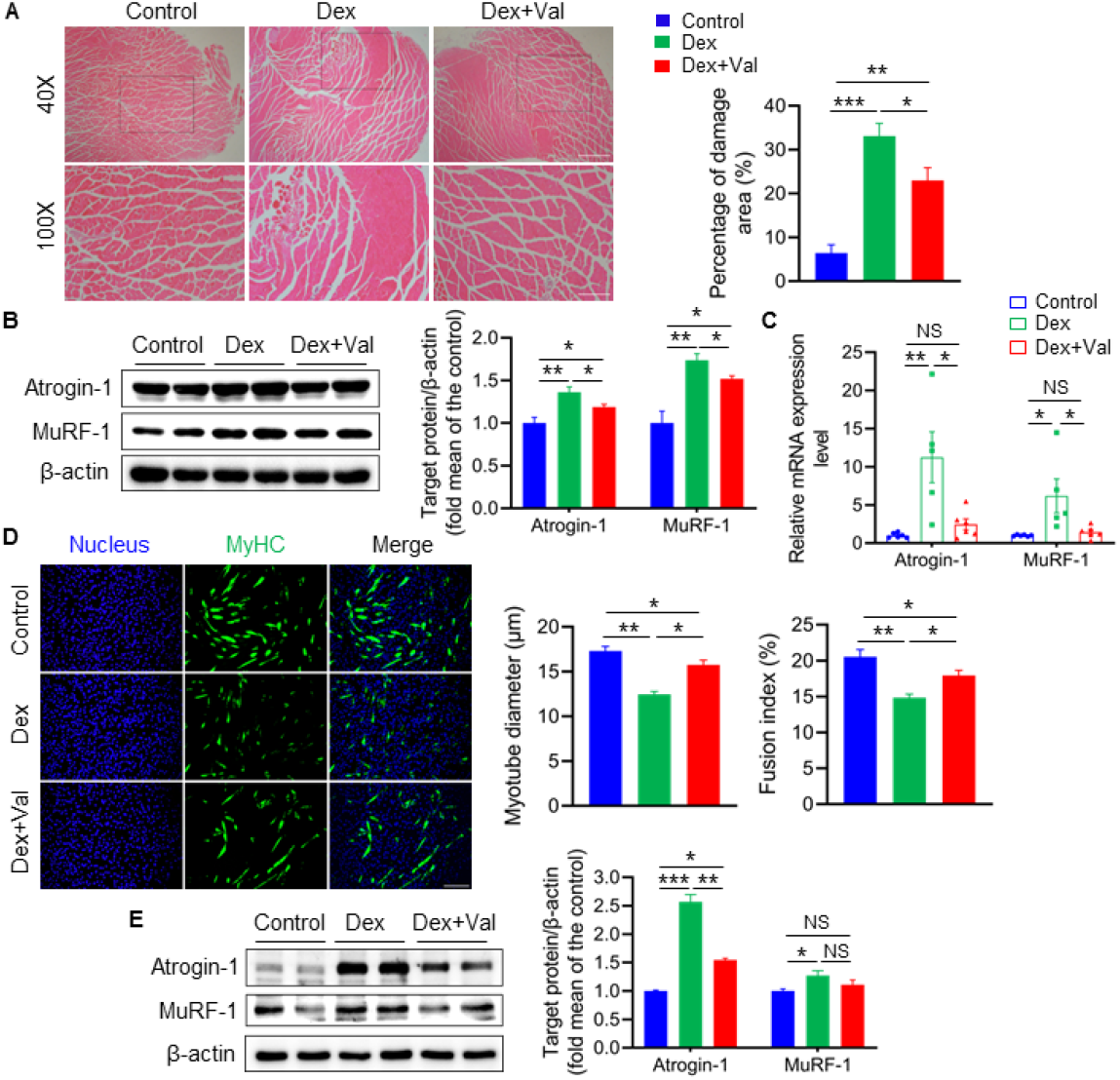
Valeric acid ameliorates Dex-induced skeletal muscle and myotube atrophy. Mice were treated with intraperitoneal injection of 20 mg/kg Dex every 2 day for two weeks and 100 mg/kg of valeric acid was fed orally every day before two weeks of Dex injection (n=5). Myotube atrophy was induced with 100 μM/L of Dex, and 5 mM/L of valeric acid was supplied at the same time (n=6). (**A**) Hematoxylin eosin staining of GA morphology. Magnification is 40×. Scale bar, 500 μm. Quantification analysis showed that Dex induced the myofiber damage, and valeric acid treatment decreased the percentage of damage area. (**B**) Western blot analysis showed that Dex induced the expression of Atrogin-1 and MuRF-1 in GA, while valeric acid treatment reduced the level of these. (**C**) Real-time PCR analysis of relative expression of atrophy genes (*Atrogin-1* and *MuRF-1*) in GA, showed valeric acid treatment could inhibit the expression of these genes induced by Dex. (**D**) Immunofluorescence stained with a specific antibody was used to identify MyHC (green) of myotube and the nucleus were stained with DAPI (blue). Magnification is 100×. Scale bar, 200 μm. Quantification analysis showed valeric acid treatment could improve the reduction of myotubes diameter and fusion index induced by Dex. (**E**) Western blot analysis showed valeric acid treatment could inhibit the expression of Atrogin-1 induced by Dex in C2C12 myotubes and had no effect on MuRF-1 induced by Dex. Statistical analysis is performed using one-way ANOVA with *Least Significant Difference test*. Data are expressed as means ± SEM. \**p* < 0.05; \*\**p* < 0.01; \*\*\**p* < 0.001; NS, not statistically significant.

## Discussion

Host genetic variations can influence the microbiota composition, and the gut microbiota can affect skeletal muscle growth and function. Here, we revealed that the gut microbiota remodeled by *MSTN* gene deletion plays a key role in regulating skeletal muscle development. *MSTN* gene knockout not only increased skeletal muscle mass, but also altered the intestinal structure and composition of intestinal flora in pigs, as shown the loss of intestinal MSTN expression, altered muscularis thickness, plica length, the increase expression of tight junction genes *ZO-1* and *Occludin*, and enriched microbioal population that produce SCFAs. We transplanted the fecal microbiota of *MSTN*^−/−^ pigs into mice, and the recipient mice had increased fast-twitch glycolytic muscle GA weight and increased levels of glycolysis proteins HK2, PFK1, and PKM2 and type IIb myofibers hypertrophy, characterized by enhanced grip strength and poor resistance to fatigue, accompanied by increased phosphorylation of the Akt/mTOR signal. Similar to the donor pigs, recipient mice were enriched in microbes that produce SCFAs. Furthermore, metabolomic analysis showed a significant increase in valeric acid levels in the colon contents. We showed that the intestinal flora remodeled by MSTN gene deletion is involved in fast-twitch glycolytic muscle growth via valeric acid, which activaties the Akt/mTOR pathway through the SCFAs receptor GPR43. Lastly, we demonstrated that valeric acid have a beneficial effect on skeletal muscle atrophy induced by Dex (Figure 8).

**Figure 8.**
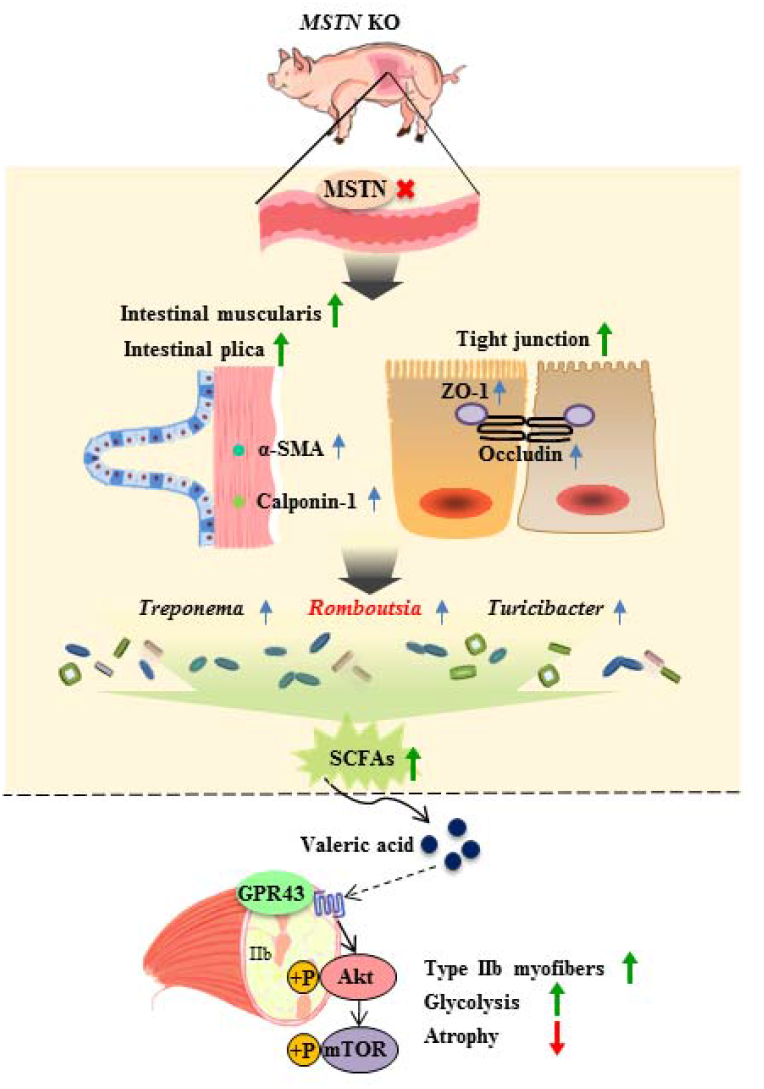
Schematic illustration. Intestine MSTN deficiency altered the intestinal structure, reshaped gut microbiota, and gut microbiota metabolite-valeric acid can activate Akt/mTOR pathway through GPR43 to stimulate fast-twitch glycolytic skeletal muscle growth.

MSTN regulates myogenic differentiation and skeletal muscle mass mainly by activating classical Smad2/3 transcription factors (Chen et al., 2021). In this study, *MSTN^−/−^* pigs generated using TALEN genome editing had significantly inhibited activation of Smad, increased CSA of type IIb myofibers, and overgrowth of skeletal muscle, which was characterized by the ‘double-muscle’ phenotype. These findings are consistent with those of the previous studies performed in *MSTN* mutant mice and cattle (McPherron et al., 1997; Ceccobelli et al., 2022; McPherron and Lee, 1997; Kambadur et al., 1997).

MSTN expression has been detected not only in the skeletal muscles but also in smooth muscles of blood vessels, penis, and other tissues and is co-localized with α-smooth muscle actin, which can affect organ functions (Verzola et al., 2017; Esposito et al., 2020; Kovanecz et al., 2017). Intestine tissues are from smooth muscle, and *MSTN* expression in intestine tissues has been confirmed; however, its role in intestine is not clear (Sundaresan et al., 2008). In this study, MSTN expression was detected in the intestine of WT pigs but not in that of *MSTN^−/−^* pigs. Importantly, this is the first study to show that *MSTN* knockout leads to a loss of its expression in the intestine and increases the intestinal muscularis thickness and plica length in pig intestine, indicating *MSTN* knockout-induced changes in intestinal morphology. The muscularis is related to intestinal motility, and its thickness can represent the ability of intestinal peristalsis; the height of the mucosal fold determines the intestinal absorption surface area (Wang et al., 2019; Zhao et al., 2017; Geda et al., 2012). In this study, increases in small intestinal muscularis thickness and plica length imply enhanced intestinal absorptive capacity in *MSTN^−/−^* pigs. Actually, *MSTN* gene mutation has also been found to affect the composition of metabolites and microbial strains in the jejunum, which might provide more useable nutrients for the host (Pei et al., 2021). The tight junction between adjacent intestinal epithelial cells is a critical component of the intestinal barrier, which provide a form of cell–cell adhesion in enterocytes and limit the paracellular transport of bacteria and/or bacterial products into the systemic circulation (Ghosh et al., 2020). Previous studies have shown that a disruption in the intestinal barrier leads to increased bacterial products of lipopolysaccharide into systemic circulation, triggering an inflammatory response in specific tissues, such as skeletal muscle or adipose tissue (Ghosh et al., 2020). On the other hand, enhancement of the intestinal barrier function effectively reduced intestinal inflammation, resulting in the alleviation of skeletal muscle loss in cancer cachexia (Sakakida et al., 2022). In this study, *MSTN^−/−^*pigs had a significantly increased expression of tight junction genes *ZO-1* and *Occludin* in intestine. It is indicated that *MSTN* gene deletion can improve the intestinal physical barrier of pigs. Importantly, changes in the intestinal environment and barrier function can alter microbial composition of the gut (Seganfredo et al., 2017; Nicoletti et al., 2017; Sekirov et al., 2010). The intestinal microflora composition of *MSTN^−/−^* pigs was analyzed, and we found that *Romboutsia*, *Treponema,* and *Turicibacter* were significantly enriched. Several studies have suggested that these microbes are involved in SCFAs production (Li et al., 2019b; Li et al., 2021; Li et al., 2019c; Bian et al., 2020). SCFAs are absorbed from the intestinal tract and play a metabolic regulatory role in different organs, which are recognized as a potential regulator of skeletal muscle metabolism and function (Frampton et al., 2020). Additionally, *Romboutsia* (Li et al., 2021; Yanni et al., 2020) and *Turicibacter* (Watanabe et al., 2021) are closely associated with metabolic disorders, such as hypertension, diabetes, dysregulation of skeletal muscle energy metabolism, and obesity. Therefore, we believe that the deletion of the *MSTN* gene in the intestine alters the intestinal structure, thus affecting the composition of intestinal flora.

FMT can transfer both host gut characteristics and metabolic phenotypes from pigs to mice (Yang et al., 2018; Diao et al., 2016; Yan et al., 2016). To verify the effect of the intestinal flora remodeled by *MSTN* gene deletion on skeletal muscle, the intestinal flora of *MSTN^−/−^* and WT pigs were transplanted into mice. Interestingly, we found that mice transplanted with *MSTN^−/−^* pigs feces had increased GA muscle mass and muscle fiber area (Figure 2). We further found that the area of type IIb myofibers increased significantly, indicating that the increased GA weight and muscle fiber area can be attributed to the growth of type IIb myofibers. Previous studies have found that Akt1 transgene activation specifically increases GA weight and type IIb myofiber growth through mTOR-dependent pathway (Izumiya et al., 2008). This is consistent with our study, where the significant activation of the Akt1/mTOR pathway was observed with increased GA mass and IIb myofiber CSA in KO-M. In addition, augment of type IIb myofibers lead to an increase in grip strength but a reduction in endurance on a treadmill test (Izumiya et al., 2008). In this study, as expected, KO-M had significantly enhanced grip strength but poor resistance to fatigue. Preservation or restoration of type IIb myofibers may delay age-related changes, mainly by reducing fat mass and liver steatosis and correcting glucose metabolism injury (Akasaki et al., 2014). Importantly, we found that SCFAs producing microbes, *Romboutsia*, enriched in *MSTN^−/−^* pigs are also significantly enriched in recipient mice (Figure 3B-D). The concentration of SCFAs was significantly increased in the colon contents. Gut microbiota transplantation from pathogen-free mice into germ-free mice was reported to increase skeletal muscle mass and reduce muscle atrophy markers, thus improving oxidative metabolic capacity. Moreover, the microbial metabolites–SCFAs treatment can also attenuate skeletal muscle impairments, especially in the GA muscle (Lahiri et al., 2019). During hibernation, the urease-producing microbes, *Alistipes*, enriched in the gut of ground squirrels can prevent muscle loss (Regan et al., 2022). These results strongly suggest that the intestinal flora remodeled by *MSTN* gene deletion is involved in the growth of fast-twitch glycolytic muscle mass and function, which may be related to the enrichment of SCFAs–producing microbes.

SCFAs are the main metabolites of intestinal microbiota and are involved in multiple physiological processes of the host (Donohoe et al., 2011; Canfora et al., 2015). Compared with untreated mice, SCFAs-treated germ-free mice showed improved muscle strength (Lahiri et al., 2019). In addition, acetic acid, a kind of SCFA, can change fiber types and regulate mitochondrial metabolism of skeletal muscle (Pan et al., 2015). Similarly, we observed that valeric acid treatment increases myotube formation in myoblasts and the skeletal muscle mass of GA in mice, especially with respect to type IIb muscle fiber formation rate. SCFAs play a downstream regulatory role mainly by binding to their receptors (Stoddart et al., 2008; Hul et al., 2019). To explore the possible mechanism of action of valeric acid in skeletal muscles, we analyzed GPRs expression and found a considerable increase in GPR43 levels following valeric acid treatment. The GPR43 receptor can activate the Akt and mTOR signaling pathway (Bian et al., 2020; Dutchak et al., 2018), which may explain the activation of the Akt signaling pathway detected in this study (Figure 6). Aging and long-term and high-dose glucocorticoid therapy could induce skeletal muscle atrophy, mainly manifesting as skeletal muscle mass loss and priority loss of type IIb muscle fibers (Akasaki et al., 2014; Haber et al., 1992; Faulkner et al., 2007; Kirkendall et al., 1998). We found that valeric acid treatment ameliorates Dex-induced myotube atrophy and partially repairs skeletal muscle atrophy (Figure 7).

In conclusion, this is the first study to demonstrate that *MSTN* gene deletion in pig intestine alters intestinal structure and function, leading to changes in the composition of intestinal microbiota. We further demonstrate that *MSTN* gene deletion-mediated remodeling of the intestinal flora increases the growth of fast-twitch glycolytic muscles. Finally, we illustrate that the microbiota metabolite valeric acid can promote myoblast differentiation and fast-twitch glycolytic myofiber growth by activating the Akt/mTOR pathway through the SCFAs receptor GPR43 and have a beneficial effect for skeletal muscle atrophy induced by Dex. These findings increase our understanding of the host genetic variation in regulating gut microbiota, and provide new insights for the treatment of muscle-related diseases, such as muscular dystrophy and sarcopenia.

## Materials and methods

### Animals

The animal study was approved by the Ethics Committee of Yanbian University (approval number SYXK2020-0009). We generated *MSTN^−/−^* pigs with 2 and 4 bp deletions in the two alleles of *MSTN* gene by TALEN genome editing technique and somatic cell nuclear transfer and these pigs were used in this experiment (Kang et al., 2017). Pigs were fed a standard commercial diet and housed in the same environmentally controlled room in a swine breeding farm. Male C57BL/6J mice aged four weeks were purchased from Vital River Laboratory Animal Technology (Beijing, China). Chow diet (Beijing HuaFuKang Bioscience, Beijing, China) and water were provided ad libitum. Mice were administered with valeric acid (100 mg/kg, Shanghai Aladdin, China) by oral gavage or water (vehicle) starting at four weeks of age, and tissues were collected after five weeks of treatment.

To establish the Dex-induced muscle atrophy model, male C57BL/6J mice aged eight weeks were intraperitoneally injected with 20 mg/kg Dex every other day for two weeks, and saline injections were used for the control group. Dex-induced skeletal muscle atrophy was determined by weight loss in mice (Hong et al., 2019; Li et al., 2017). A total of 100 mg/kg of valeric acid was provided orally to mice every day two weeks before Dex injection until the full experiment cycle. Mice were raised in a pathogen free environment at a controlled ambient 21±1 L, 40–60% relative humidity, and 12/12 h cycle of alternating day and night. In all experiments, the animals were fasted overnight before they were euthanized.

### Fecal microbiota transplantation

Fecal samples were collected daily from six-month old *MSTN^−/−^*and wild-type (WT) donor pigs in the morning. In a sterile environment, they were homogenized and suspended using sterile saline (250 mg/mL), and the mixture was centrifuged at 800 × g for 5 min. Antibiotics mixture (50 μg/mL streptomycin, 100 U/mL penicillin, 170 μg/mL gentamycin, 100 ug/mL metronidazole, 125 ug/mL ciprofloxacin; all from Sigma) was added to sterile drinking water and was given daily for one week before FMT. From five weeks of age, each group of recipient mouse was gavaged with 200 μL of the corresponding bacterial suspension every day for eight weeks until tissue collection.

### Analysis of gut microbiota

The fecal samples used for microbiota analysis were collected separately from donor pigs at six-month old and recipient mice after eight weeks of FMT. Methods used to analyze the diversity and taxonomic profiles of gut microbiota in donor pigs and recipient mice have been described previously (Quan et al., 2020). Briefly, the CTAB method was used extract the total genomic DNA from fecal bacteria. DNA sample with a final concentration of 1 ng/μL was used for bacterial 16s rRNA gene amplification seuqnecing (V3-V4 regions). The Illumina NovaSeq platform (Novogene, Beijing, China) was used to determine the sequencing abundance and diversity of the intestinal flora in pigs and mice. The library quality was assessed using a Qubit@ 2.0 Fluorometer (Thermo Scientific) and Agilent Bioanalyzer 2100 system.

Paired-end reads were allocated according to the unique barcodes of the sample and truncated by cutting off the barcode and primer sequences. FLASH (v1.2.7) was used to merge the overlapped reads between paired-end reads. According to the QIIME (V1.9.1) quality control process, high-quality clean tags were obtained from qualitative filtration of the original reads under specific filtration conditions. The effective tags were finally collected by comparing the sample tags with the reference database (Silva database) after the detection and removal of chimera sequences using the UCHIME algorithm. The QIIME software was used to calculate all indices in the samples, and R (v2.15.3) was used for bioinformatic analyses of the sequences. The same operational taxonomic units had at least 97% similarity in sequences. Alpha diversity, beta diversity, and principal component analysis (PCA) were described according to the unweighted unifrac distances.

### Cell culture

C2C12 myoblasts (1 × 10^5^ cells/well) were cultured in six-well culture plates in Dulbecco’s modified Eagle’s medium (DMEM; Invitrogen-Gibco), containing 10% fetal bovine serum (Sigma), 100 U/mL penicillin and 100 U/mL streptomycin (Invitrogen-Gibco) for proliferation. For differentiation, C2C12 myoblasts at 80% confluence were induced to differentiate in DMEM with 2% horse serum (Invitrogen); valeric acid and isobutyric acid were added to the differentiation medium for 24 h. Cells were supplemented with a fresh differentiation medium every two days. Myotubes were obtained on day 5 of the differentiation phase.

To establish the Dex-induced myotube atrophy model, myoblasts were treated with 100 μm/L of Dex at the beginning of differentiation for 24 h, and 5 mM/L valeric acid was added to the treatment group. Myoblasts were cultured in a fresh differentiation medium for five days. Myotubes were stained with anti-MyHC antibody (MyHC, A4.1025, Sigma), and Alexa Fluor 488-labelled goat anti-mouse IgG was used as secondary antibody (Jackson ImmunoResearch Laboratories). The nuclei were counterstained with 10 μg/μL DAPI (D-9106, Beijing Bioss Biotechnology). The diameter and the number of nuclei of the differentiated myotubes were measured using Image J (1.51q, National Institutes of Health, USA). For each group, five pictures were randomly taken from each well of the six-well plates. The diameters of the three different parts of each myotube were measured, and the average value was calculated. To determine the C2C12 fusion index, the number of nuclei in the myotubes were calculated and divided by the total number of nuclei and multiplied by 100.

### Histological analysis

Skeletal muscle and intestine morphology were detected by hematoxylin eosin (HE) staining. The longissimus dorsi and intestine tissues from *MSTN^−/−^* and WT pigs were from 5 µm thick paraffin-embedded sections. Images were obtained using a light microscope (BX53, Olympus, Japan).

Liquid nitrogen-cooled isopentane was used to rapid freeze the skeletal muscle tissues, which were embedded in the OCT compound (Sakura Finetech USA Inc.). Cryostat sections (10 µm) were prepared from the mid-belly of the muscle tissue. Fiber-type of skeletal muscle was stained using MyHC type I (BA-D5, DSHB, Douglas Houston), MyHC type IIa (SC-71, DSHB, Douglas Houston), MyHC type IIb (BF-F3, DSHB, Douglas Houston), and laminin (ab11575, Abcam) monoclonal antibodies. Alexa Fluor 647 conjugate goat anti-mouse IgG2b, Alexa Fluor 488 conjugate goat anti-mouse IgG1, Alexa Fluor 555 conjugate goat anti-mouse IgM, or Alexa Fluor 594 conjugate goat anti-rabbit IgG were used as secondary antibodies. The nuclei were counterstained with 10 μg/μL of DAPI. Fluorescence was detected using a confocal laser scanning microscope (FV3000, Olympus, Japan). Image J software was used to measure the thickness and the cross-sectional area (CSA) of the myofibers. The damage area of skeletal muscle fiber was evaluated by calculating the ratio of muscle fiber ablation area to the total muscle fiber CSA.

### Quantitative real-time PCR

Total RNA was extracted from liquid nitrogen quick-frozen tissue using a Total RNA Extraction Kit (LS1040; Promega) as per the manufacturer’s protocol. After evaluating the concentration and purity of RNA, an equal amount of RNA was used for reverse transcription. Information on the primers used is available in supplementary materials. Real-time PCR was performed on Mx3005P system (Agilent, Santa Clara, CA, USA), and the relative gene expression levels were calculated using the 2^-△△CT^ method and normalized to that of the control group.

### Western blotting

The cells and tissues were homogenized in RIPA buffer (Beyotime). Equal amounts of protein were calculated and loaded according to the concentration of protein that was detected by a BCA kit (Beyotime, Shanghai, China). Subsequently, immunoblot analysis was performed following standard procedures. Protein samples were electrophoresed, transferred, blocked and incubated with the following primary antibodies: phospho-Akt (Ser473), phospho-mTOR (Ser2448), phospho-Smad2 (Ser465/467)/Smad3 (Ser423/425), Akt, mTOR, Smad2/3, and HK2 from Cell Signaling Technology; PFK1 and PKM2 from Shanghai Absin, Inc.; MyHC type I, MyHC type Iia, and MyHC type IIb from DSHB; MSTN, MyoD, MyoG, α-SMA, calponin-1, MuRF-1, atrogin-1, actin, and tubulin from Beijing Bioss Biotechnology, Inc.. The ChemiDoc™ MP Imaging System and Image Lab software (Bio-Rad, Shanghai, China) were used to analyze the blot bands.

### SCFAs analysis

Colon contents were collected at the time of mouse tissue sample collection after eight weeks of FMT. SCFAs were extracted from mouse feces using 1:1 acetonitrile:water solution and derivatized using 3-nitrophenylhdyrazones. SCFAs were analyzed using a Jasper HPLC coupled to a Sciex 4500 MD system (LipidALL Technologies Co., Ltd, Changzhou, China). In brief, a Phenomenex Kinetex C_18_ column (100 × 2.1 mm, 2.6 µm) was used to separate individual SCFA. The mobile phase A consisted of 0.1% formic acid aqueous solution, and the mobile phase B consisted of 0.1% formic acid acetonitrile. Octanoic acid-1-^13^C_1_ (Sigma-Aldrich) and butyric-2,2-d_2_ (CDN Isotopes) were used as internal standards for quantitation (Li et al., 2019a).

### Statistical analysis

Statistical analysis was performed using SPSS (17.0, IBM, Armonk, NY, USA) and GraphPad Prism (San Diego, CA, USA). Data are presented as the mean ± SEM, and were compared using a repeated measure two-way analysis of variance (ANOVA), one-way ANOVA, or Student’s *t-test*. Statistical significance was set at \**P*<0.05, \*\**P*<0.01, \*\*\**P*<0.001.

## Acknowledgments

The author would like to appreciate Yanbian University for its support to Tumen River Scholars.

## Additional information Competing interests

The authors declare that they have no competing interests.

## Author contributions

L-HQ, J-DK, Z-BL and X-JY conceived the project, contributed to experimental design; L-HQ, J-DK, Z-BL and SH performed experiments, interpreted the results, prepared the figures and wrote the manuscript; SH, HL and H-LL contributed to writing and editing; JW, CH, DW, YL and DL performed animal studies; Z-BL, KG and BQ performed animal tissue analysis; SC and MX performed cellular experiments; all authors discussed the results and approved the manuscript.

## Ethics approval and consent to participate

The animal study was approved by the Ethics Committee of Yanbian University (approval number SYXK2020-0009).

## Additional files

### Supporting information

**Supplementary Figures and Table.**

Supplementary methods.

## Availability of data and materials

The raw reads of 16s rRNA gene sequences have been submitted to the NCBI BioSample database (Porcine data: PRJNA743164; Mice data: PRJNA743401).

All sample metadata and intermediate analysis files are available at https://www.scidb.cn/s/YRZ32q.

