## Supplement information for "Myostatin gene deletion alters gut microbiota stimulating fast-twitch glycolytic muscle growth"

*Corresponding author

**Additional file 1: Figures S1-S4.**

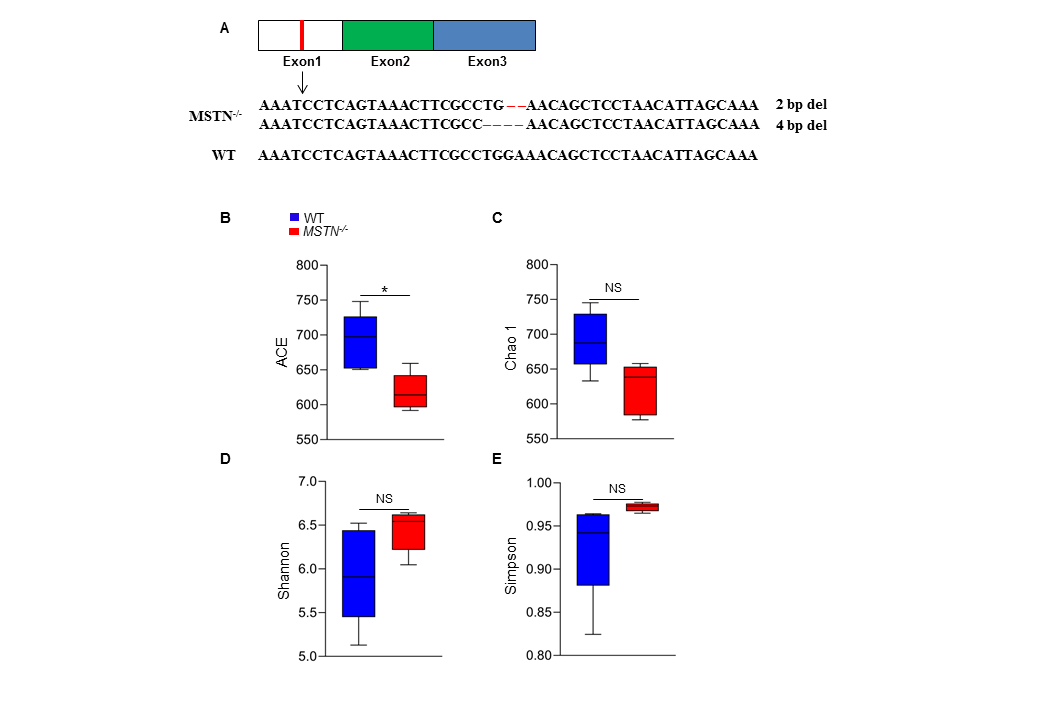

Figure S1. Gene sequence and alpha-diversity of microbiota of *MSTN^−/−^* pigs. (A) Gene sequence of *MSTN^−/−^* pigs generated by genome editing. (B) Alpha-diversity analyses showed that the ACE was lower in *MSTN^−/−^* pigs and had no difference in (C) Chao 1, (D) Shannon, (E) Simpson index between *MSTN^−/−^* and WT pigs. Statistical analysis is performed using Student’s *t-test* between WT and *MSTN^−/−^* pigs. Data are means ± SEM. **p* < 0.05; NS, not statistically significant.

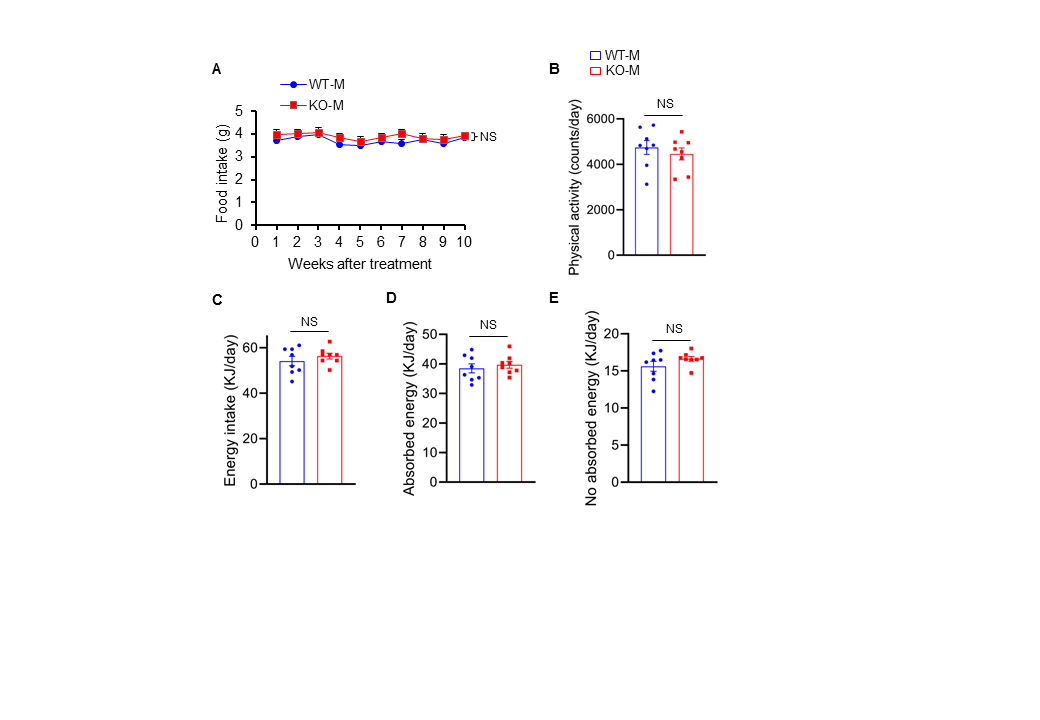

**Figure S2. Food intake, physical activity and energy in mice after fecal microbiota transplantation.** Mice were treated with porcine fecal microbiota for 8 weeks by daily oral gavage after combined antibiotics treatment for a week. WT-M, WT pigs fecal microbiota-received mice (n=8); KO-M, *MSTN^−/−^* pigs fecal microbiota-received mice (n=8). There were no differences between WT-M and KO-M in (**A**) Food intake, (**B**) physical activity, (**C**) energy intake (**D**) absorbed energy, (**E**) no absorbed energy. For food intake curves, a repeated measure two-way ANOVA, others is performed using Student’s *t-test* between WT-M and KO-M groups. Data are means ± SEM. NS, not statistically significant.

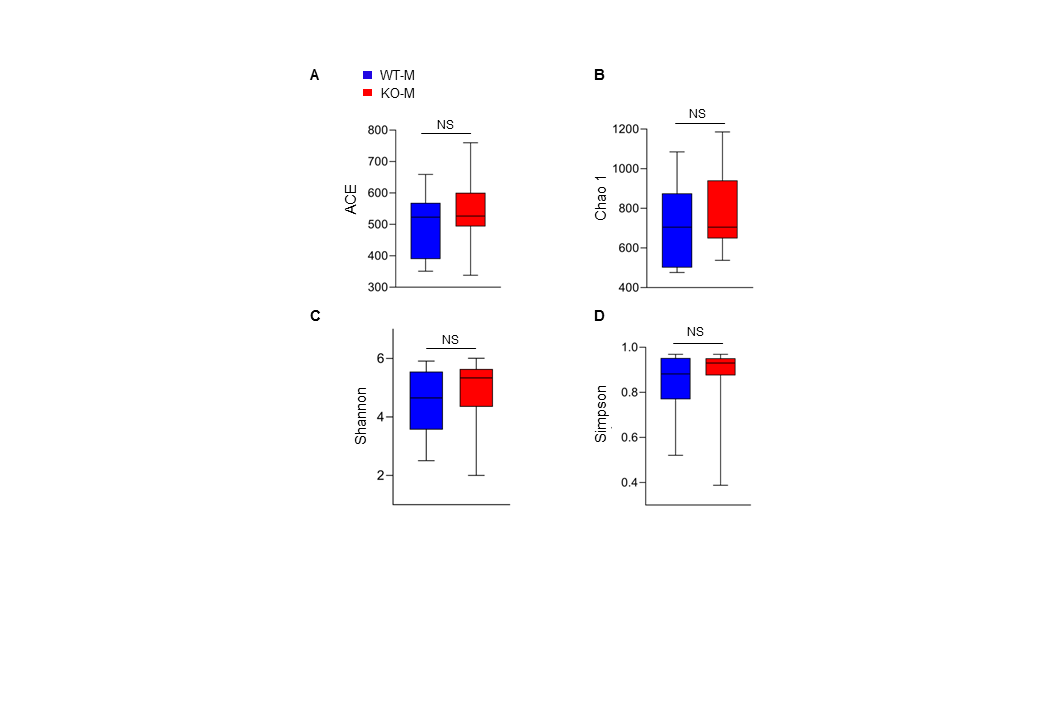

**Figure S3. Alpha-diversity in mice after fecal microbiota transplantation from *MSTN^−/−^* pigs.**

Mice were treated with porcine fecal microbiota for 8 weeks by daily oral gavage after combined antibiotics treatment for a week. WT-M, WT pigs fecal microbiota-received mice (n=8); KO-M, *MSTN^−/−^* pigs fecal microbiota-received mice (n=8). There were no differences between WT-M and KO-M in (**A**) ACE, (**B**) Chao 1, (**C**) Shannon and (**D**) Simpson index. Statistical analysis is performed using Student’s *t-test* between WT-M and KO-M groups. Data are means ± SEM. NS, not statistically significant.

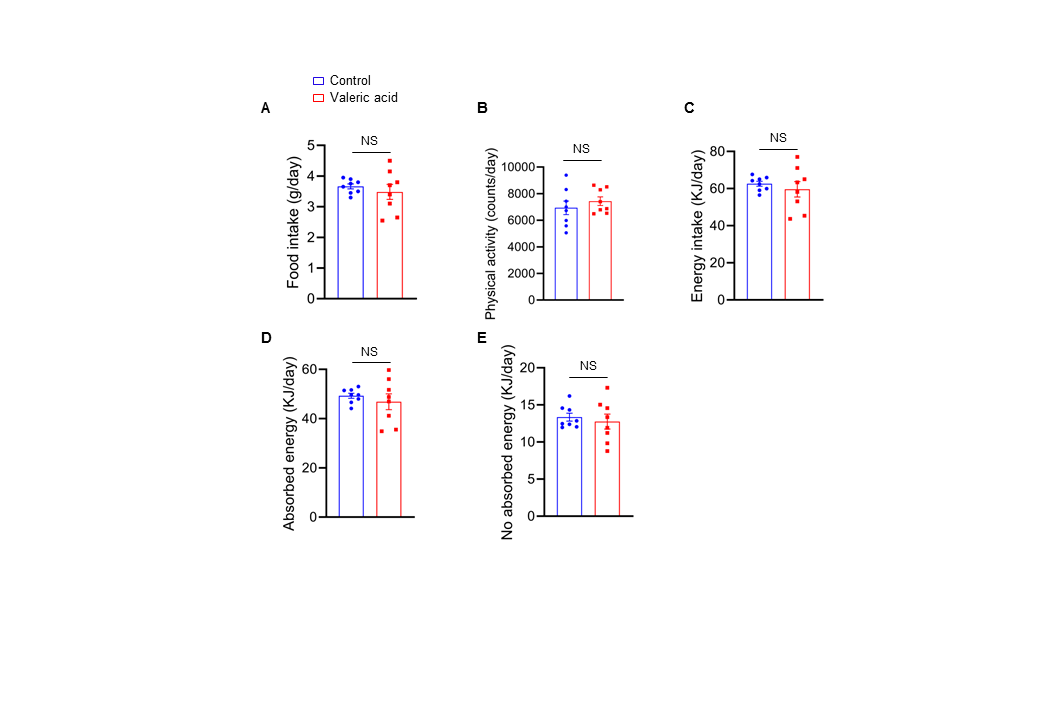

**Figure S4. Food intake, physical activity and energy in mice after valeric acid treatment.** Mice were treated with valeric acid (100 mg/kg) for five weeks by daily oral gavage (n=8). There were no differences between valeric acid treatment and control in (**A**) Food intake, (**B**) Physical activity, (**C**) Energy intake (**D**) Absorbed energy, (**E**) No absorbed energy. Statistical analysis is performed using Student’s *t-test*. Data are means ± SEM. **p* < 0.05; NS, not statistically significant.

Table S1.

Primers sequences used for real-time PCR

| Gene | Primer sequence (5’-3’) | GeneBank accession number |
| --- | --- | --- |
| ZO-1 | Forward: AAGCCCTAAGTTCAATCACAATCT | XM_021098896.1 |
|  | Reverse: ATCAAACTCAGGAGGCGGC |  |
| Occludin | Forward: TCCTGGGTGTGATGGTGTTC | XM_005672525.3 |
|  | Reverse: CGTAGAGTCCAGTCACCGCA |  |
| Claudin-1 | Forward: AAACCGTGTGGGAACAACCA | NM_001244539.1 |
|  | Reverse: CACATGAAAATGGCTTCCCTC |  |
| GAPDH | Forward: GCCATCACCATCTTCCAGG | AF017079 |
|  | Reverse: TCACGCCCATCACAAACAT |  |
| GPR43 | Forward: ATGTAGCCGATGGAAGGAAGAG | NM_146187.4 |
|  | Reverse: CAGCCCATGTCTTCACGGT |  |
| GPR41 | Forward: TCTTGTATCGACCCCCTGGT | NM_001033316.2 |
|  | Reverse: CTCCTGCGGTCCACTCTTTT |  |
| GPR109a | Forward: GTGAACCCGATAATAACCGAAGC | NM_030701.3 |
|  | Reverse: TGAGCGGCAATCTCCCTTCT |  |
| Atrogin-1 | Forward: GACTGGACTTCTCGACTGCC | NM_026346.3 |
|  | Reverse: TCAGGGATGTGAGCTGTGAC |  |
| MuRF-1 | Forward: GAGGGGCTACCTTCCTCTCA | NM_001039048.2 |
|  | Reverse: CCAGAGCGTGTCTCACTCAT |  |
| β-actin | Forward: GTACCACCATGTACCCAGGC | NM_007393.5 |
|  | Reverse: AACGCAGCTCAGTAACAGTC |  |

**Additional file 2: Supplementary methods.**

**Wheel running and grip strength**

Before the wheel running experiment, all mice were trained to run aton at a low speed for a week. Mice were individually housed in running wheels (SA102, Jiangsu SANS Bioscience, China), according to the manufacturer's recommendations. Briefly, the mice ran on the wheel at a speed of 30 r/min until they dropped, and the time and distance they ran were recorded. The mice were trained for a week to use their limbs to grasp a grip meter (SA417, Jiangsu SANS Bioscience, China) before the grip strength test. Each mouse was tested for the highest peak strength by making them pull the grip dynamometer horizontally.

**Physical activity and energy metabolic**

Each mouse was individually measured by a small animal activity detector according to the manufacturer's instructions (SA-YLS-1C, Jiangsu SANS Bioscience, China). The physical activity of mice was measured in a quiet environment for 48 h. Food intake and fecal output were collected and weighed, and food loss was subtracted.

The collected mouse feces were dried at 65℃ for 24 h, and the calorific value in dry feces and food was accurately measured in isothermal 22℃ by oxygen bomb calorimeter (IKA^®^c2000 basic, Germany). Each mouse's daily energy intake, energy absorbed, energy expelled and energy unabsorbed were calculated by caloric.

**Free fatty acids (FFAs) analysis**

FFAs were extracted from mouse feces using a modified version of the Bligh and Dyer’s method (LipidALL Technologies Co., Ltd, Changzhou, China). Briefly, fecal samples were homogenized with 750 µL of chloroform: methanol 1:2 (v/v) and 10% deionized water, and incubated at 4℃ for 30 min. The samples were centrifuged after addition of 250 µL of chloroform and 350 µL of deionized water. Lipid in the lower organic phase after centrifugation was extracted twice. After that, the total extract was collected and dried in the SpeedVac under OH mode.

Agilent 1290 UPLC combined with a triple quadrupole/ion trap mass spectrometer (6500 Plus Qtrap; SCIEX) was used for FFAs analysis. Normal phase (NP)-HPLC with a Phenomenex Luna 3 µm-silica column (internal diameter 150 × 2.0 mm) was used for lipids separation. The conditions as follows; chloroform: methanol: ammonium hydroxide (89.5:10:0.5) was used as mobile phase A, and chloroform: methanol: ammonium hydroxide: water (55:39:0.5:5.5) was used as mobile phase B. D31-16:0 (Sigma-Aldrich) and d8-20:4 (Cayman Chemicals) was used as internal standards for FFAs quantitation.
